# Tomato Prosystemin is much more than a simple Systemin precursor

**DOI:** 10.1101/2021.09.15.460427

**Authors:** Donata Molisso, Mariangela Coppola, Martina Buonanno, Ilaria Di Lelio, Simona Maria Monti, Chiara Melchiorre, Angela Amoresano, Giandomenico Corrado, John Paul Delano Frier, Andrea Becchimanzi, Francesco Pennacchio, Rosa Rao

## Abstract

Systemin (Sys) is an octadecapeptide which, upon wounding, is released from the carboxy terminus of its precursor, prosystemin(ProSys) to promote plant defenses. Recent findings on the disordered structure of ProSysprompted us to investigate a putative biological role of the whole precursor deprived of Sys peptide. We produced transgenic tomato plants expressing a truncated *ProSys* gene in which the exon coding for Sys was removed and compared their defense response with that induced by the exogenous application of the recombinant deleted ProSys[ProSys_(1-178)_].By combining protein structure analyses, transcriptomic analysis, gene expression profiling and bioassays with different pests we demonstrate that the truncated ProSys, that does not induce the endogenous *ProSys* gene, promotes defense barriers in tomato plants through a hormone independent defense pathway, likely associated with the production of oligogalacturonides (OGs). Both transgenic and plants treated with the recombinant protein showed the modulation of the expression of genes linked with defense responses and resulted protected against the lepidopteran pest *Spodoptera littoralis* and the fungus *Botrytis cinerea*. Our results suggest that the overall function of the wild type prosystemin is more complex than previously shown as it might activate at least two tomato defense pathways: the well-known Sys-dependent pathway connected with the induction of JA biosynthesis and the successive activation of a set of defense-related genes and the ProSys_(1-178)_-dependent pathway associated with OGs production leading to the OGs mediate plant immunity.

## Introduction

The in-depth studies on plant defense responses in *Solanaceae* have shown the important role of signaling peptides associated with tissue wounding and insect herbivory (McGurl *et al*., 1994; Pearce *et al*., 1991; Ryan, 2000). These peptides include tomato Systemin (Sys), an octadecapeptide released upon wounding from prosystemin (ProSys), a precursor protein of 200 amino acid, through a poorly defined pathway, apparently mediated by phytaspases(Beloshistov *et al*., 2018; McGurl *et al*., 1992). Sys promotes long-distance defense responses by amplifying the jasmonate signaling pathway, which appears to be central to systemic defense signaling (Schilmiller and Howe, 2005). Other members of the same family of defensive peptides are hydroxyproline-rich systemins (HypSys), which are also released from a larger precursor (Pearce *et al*., 2008). Although these peptides are structurally unrelated to Sys, they took their name due to their systemin-like function (Pearce, 2011). ProSys and HypSys work cooperatively in the regulation of tomato defense responses (Narváez-Vásquez *et al*., 2007).

The modulation of direct and indirect defenses against herbivorous insects by Sys has been widely characterized (Corrado *et al*., 2007; Degenhardt *et al*., 2010; Ryan, 2000). The constitutive expression of the *ProSys* gene in tomato plants triggers the increase of protease inhibitors (PIs) and other defensive compounds conferring resistance to chewing and sucking insects, phytopathogenic fungi and salt stress(Coppola *et al*., 2015; Dombrowski, 2003; McGurl *et al*., 1992; McGurl *et al*., 1994; Orsini *et al*., 2010; Ryan, 2000; Zhang *et al*., 2020). In addition, transgenic plants are characterized by an increased level of indirect defense barriers compared to untransformed controls, showing a higher level of attractiveness towards natural enemies of phytophagous insects(Corrado *et al*., 2007; Degenhardt *et al*., 2010; El Oirdi *et al*., 2011). This is probably the consequence of the deep transcriptomic reprogramming observed in transgenics, resulting in more than 500 differentially expressed genes (Coppola *et al*., 2015). The analysis of these genes showed that *ProSys* overexpression, on one hand, reduces the expression of genes related to carbohydrate metabolism and, on the other, promotes the expression of a series of defense genes regulated by different signaling pathways, thus cross-modulating growth and defense pathways. The Sys peptide is considered to be solely responsible for the biological activity in tomato, as suggested by the increased production of protease inhibitors (PI) upon Sys application on wounded or intact stems and leaves that didn’t occur when a truncated Sys-less version of ProSys was used (Coppola *et al*., 2019; Dombrowski *et al*., 1999; Pearce *et al*., 1991; Ryan and Pearce, 2003; Schilmiller and Howe, 2005). Thus, a single peptide appears to trigger multiple defense pathways in response to a wide range of stress agents (Corrado *et al*., 2007).

The mechanism underlying such a large ‘anti-stress’ capacity, associated with a single peptide, is difficult to understand from a functional point of view. Perhaps Sys ‘is not alone’ in the activation of tomato defense responses as suggested by the structural features of ProSys protein. In fact, it was very recently demonstrated that ProSys is an intrinsically disordered (ID) protein(Buonanno *et al*., 2018) without a stable or ordered three-dimensional structure. ID proteins (IDPs) play a central role in regulating the transduction pathways of various signals, including the ‘defense signal’ in addition to other crucial cellular processes, such as the regulation of transcription and translation (Oldfield *et al*., 2008). It has been proposed that the plasticity of IDPs helps sessile organisms, like plants, in establishing complex networks in response to the exposure to a myriad of both biotic and abiotic stress agents, from which they cannot move away to prevent damage(Hamdi *et al*., 2017). The characteristic flexibility of IDPs allows them to assume a number of conformations able to target multiple molecular partners (Tompa, 2002). Therefore, based upon above assumptions, we postulate that ProSys can be more than a simple precursor of Sys, and that it integrates additional functions likelyactivating multiple stress-related pathways upon interacting with different molecular partners. This hypothesis is also supported by the altered proteomic profile and increased resistance against *Botrytis cinerea* observed in tobacco transgenic plants constitutively expressing the truncated *ProSys* which lacks the Sys-encoding exon (Corrado *et al*., 2016).

Here we provideseveral evidences in support of this intriguinghypothesis. A wealth of molecular and functional data ontransgenic tomato plants, constitutively expressing the truncated *ProSys*cDNA, and on their interactions with *S. littoralis*and *B. cinerea*, have been gathered, showing a multifaceted enhanced resistance against these biotic stress agents. These results have been duplicated by exogenous treatments with a recombinant ProSys deprived of the Sys region (hereafter referred as ProSys_(1-178)_protein), which provides further direct evidence in support of the multifunctional role of ProSys in orchestrating the defense response of tomato plants.

## Results

### Production and characterization of transgenic tomato plants expressing ProSys_(1-178)_

Tomato plants were stably transformed via *Agrobacterium tumefaciens* with the pPRO8 vector (Corrado *et al*., 2016), carrying a construct containing a ProSys cDNA sequence lacking the region coding for the Sys peptide. A schematic representation of the transgenic T-DNA is shown in FigureS1a. The truncated ProSys was under control of the constitutive CaMV 35SRNA promoter and the pea *rbcs* terminator (FigureS1a). Putative transformants were screened by PCR (FigureS1b). As expected, genotypes showing different transgene expression levels were obtained (FigureS1c). Transgenic genotypes homozygous for a single-copy T-DNA insertion and high levels of transgene expression were selected by growing plants up to T_4_ generations on kanamycin enriched media and analyzing progenies for kanamycin resistance and PCR. Two homozygous lines (indicated as line 1 and line 2) were selected for further investigations.

In this lines, the amplification of a specific region of the endogenous *ProSys* gene by Real Time RT-PCR, using primers annealing on *ProSys* 3’-UTR, (Figure1a, 1b, Table S1) allowed to establish that the truncated ProSys did not influence the expression of endogenous *ProSys* gene (Figure1c). This result indicated the suitability of the transgenic plants produced to evaluate the effect of truncated *ProSys* gene on tomato defense responses.

**Figure1:**
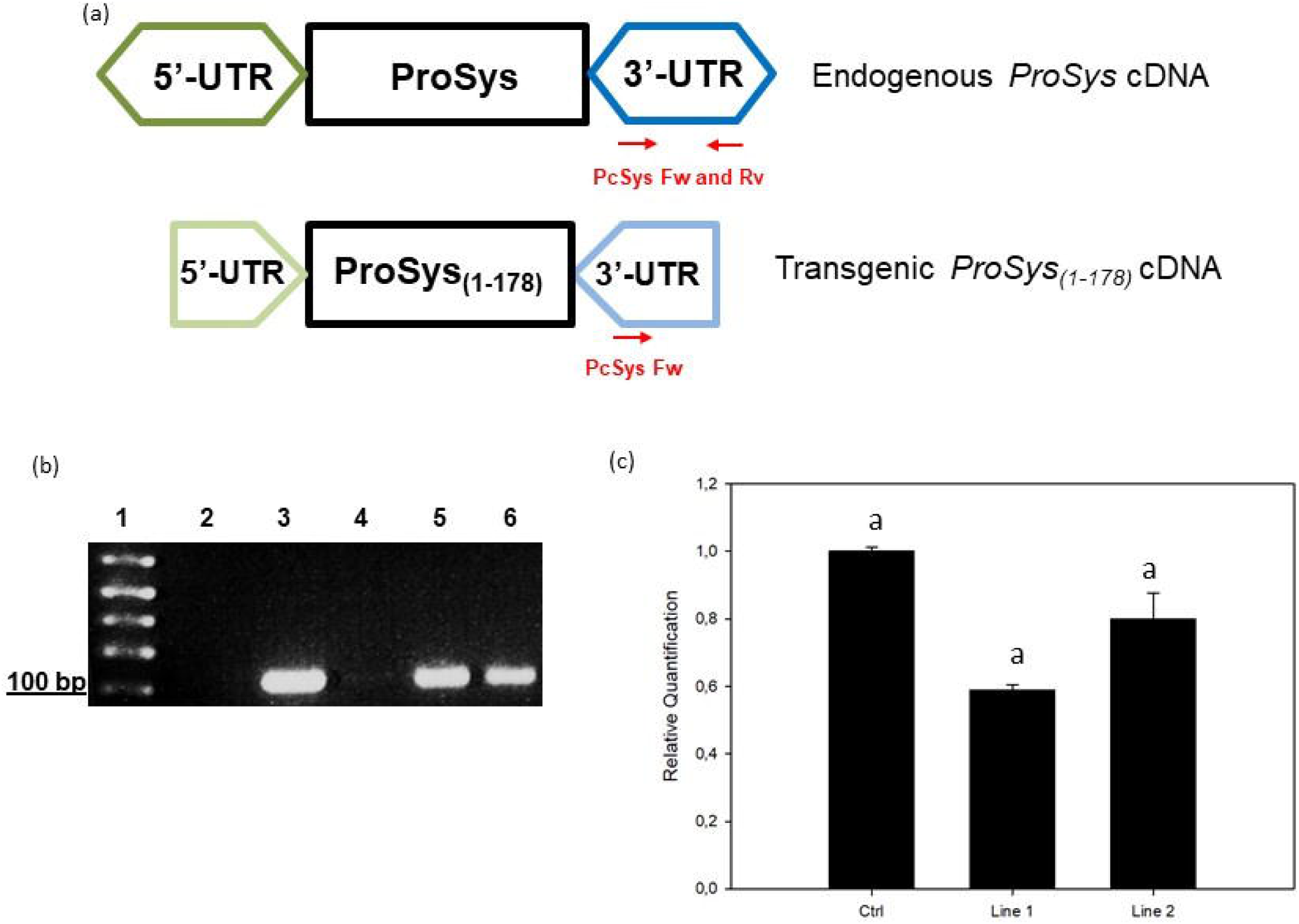
Expression analysis of endogenous *ProSys*. **a**) PCR amplification strategy of endogenous *ProSys*gene.Primers couple named PcSysFw and Rv allows to discriminate the endogenous *ProSys* from the transgenic expression cassette due to its annealing on the 3’-UTR region (dark blue hexagon); PcSys reverse primer annealing is impaired on the transgenic cassette due to the truncation of 3’-UTR (light blue pentagon). (**b**) RT-PCR of endogenous ProSys on transgenic and control plants. Lane 1: 1 kb Plus ladder (Thermo Fisher Scientific); lane 2: control (no template); lane 3: PCR positive control; lane 4: amplification of ProSys cDNA in untrasformed plants; lane 5-6: amplification of ProSys_(1-178)_line 1 and 2 with PcSys primers. (**c**) Relative quantification (RQ) of the endogenous prosystemin gene expression by Real-time RT-PCR. Amplification has been carried out using primers annealing on 3’-UTR region (absent in transgenic expression cassette). RQ are shown relative to the calibrator genotype Red Setter. No significant differences in gene expression levels were detected between transgenic and control plants.

### Production and characterization of the biochemical features of ProSys_(1-178)_

cDNA encoding ProSys_(1-178)_ was cloned in the pETM11 expression vector designed to obtain a protein having a N-terminal histidine tag. The recombinant protein was expressed in *E. coli* BL21(DE3) strain and was highly purified after two purification steps with a final yield of 1 mg/l culture. Protein identity was confirmed by LC-ESI-MS analysis. As expected, ProSys_(1-178)_ showed the peculiar features of an IDP, as observed for the full length protein (Buonanno *et al*., 2018). In particular, ProSys_(1-178)_ migrated as a protein with a greater molecular weight on SDS-PAGE and eluted as a protein with an apparent molecular mass of 54.4 kDa by SEC (FigureS2a,S1b). Light scattering analysis confirmed that, in solution, ProSys_(1-178)_ is a monomer with a molecular weight of 21.1±0.2 kDa (FigureS2c), thus highlighting its scarce compactness as already reported for the full-length protein. Indeed, CD spectra revealed a disorder content (Figure2), a reversible temperature-induced behavior, and a capability to increase secondary structure content in the presence of TFE co-solvent (Figure S3a,S3b). The presence of an isodichroic point at 203.5 was consistent with a coil-helix transition.

**Figure2:**
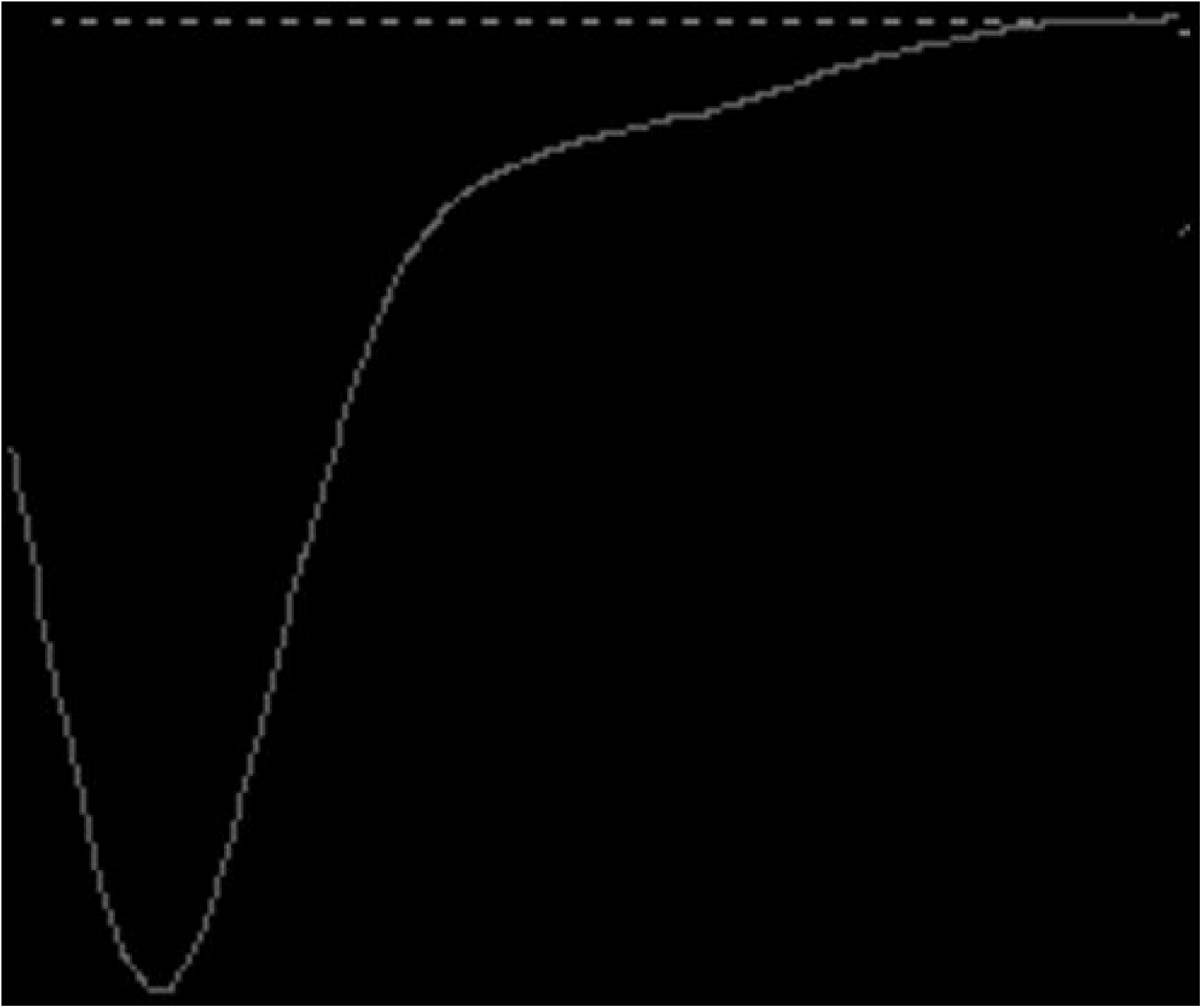
ProSys_(1-178)_recombinant proteinis intrinsically disordered. TheFar-UV circular dichroism spectrum was recorded from 260 nm to 190 nm at 20°C in 10 mM phosphate buffer pH 7.4 at a protein concentration of 6.8 μM.

### ProSys_(1-178)_ enhances plant resistance against *S. littoralis*and *B. cinerea*

#### Transgenic plants assays

*S. littoralis* larvae fed with ProSys_(1-178)_transgenic leaf disks were severely impaired in their growth (Figure3a) and showed higher mortality rates compared to controls (Log-Rank test: χ2 = 21.19, df = 3, *P*< 0.0001) (Figure3b). From day 5 until pupation, larval weights were significantly lower for larvae fed on the 2 transgenic lines than on controls (Figure3a, Table S2) (One Way ANOVA: *P*< 0.0001).

**Figure3:**
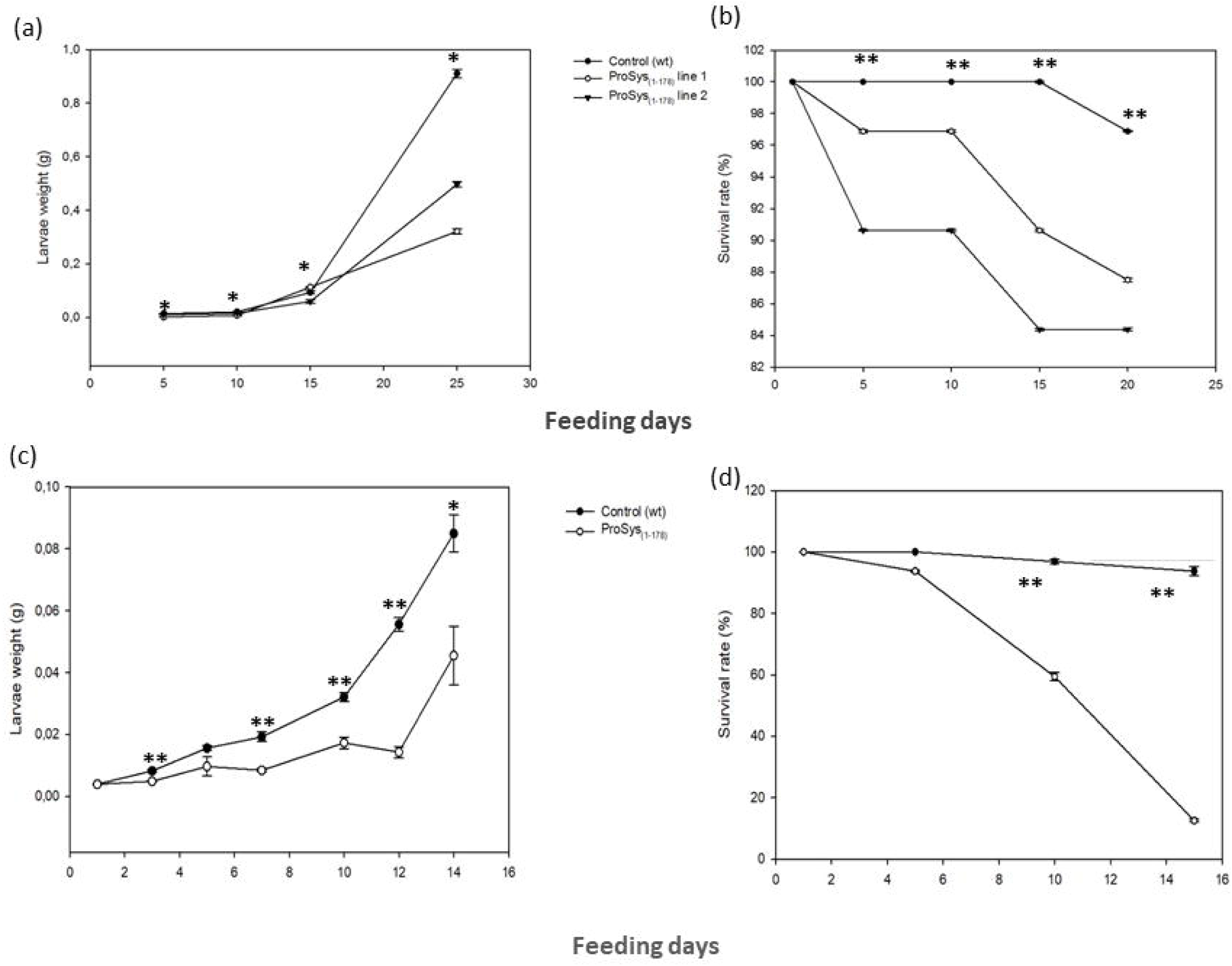
Enhanced resistance of ProSys_(1-178)_ transgenic lines to *S*.*littoralis* larvae (a,b) and of leaves of tomato plants exogenously treated with ProSys_(1-178)_recombinant protein (c,d). (**a**,**d**) Larval weight increase upon feeding with leaves from ProSys_(1-178)_ transgenic lines (ProSys_(1-178)_ lines 1 and 2 and control plants and upon upon feeding with leaves from ProSys_(1-178)_-treated and control plants. The graphs display the average (± S.D.) of larval weight at several feeding days (**a**, * One-way ANOVA: *P*< 0.0001; **b**, Student’s t-test: *\* P* <0.01; *\*\* P* <0.0001). (**b**,**d**) Survival rate of larvae fed on leaves transgenic or control plants and on ProSys_(1-178)_-treated and control leaves (** Log-Rank test: *P*< 0.0001). Asterisks denote mean values that are significantly different.

The transgene effect caused a strong reduction of *B. cinerea* colonization with the consequent reduction of necrosis areas (Figure4a).

#### Plant treatments with exogenous ProSys_(1-178)_

The exogenous application of recombinant protein ProSys_(1-178)_duplicated the enhancement of defense barriers observed with transgenic plants, confirming a direct role of the protein in triggering plant defense responses. Indeed, *S. littoralis*larvae fed with ProSys_(1-178)_-treated leaves showed, compared to controls, a significant reduction of their weight starting from day 3until day 14 (Student’s t-test: *P*< 0.0001) (Figure3c, Table S3), and a higher mortality rate, which reached 100% by day 15(Log-Rank test: χ2= 59,75, df = 1, *P*< 0.0001) (Figure3d).

The effect of the plant treatment with the recombinant ProSys_(1-178)_ on *B. cinerea* resulted in a significant reduction of lesion areas (Figure 4b).

**Figure4:**
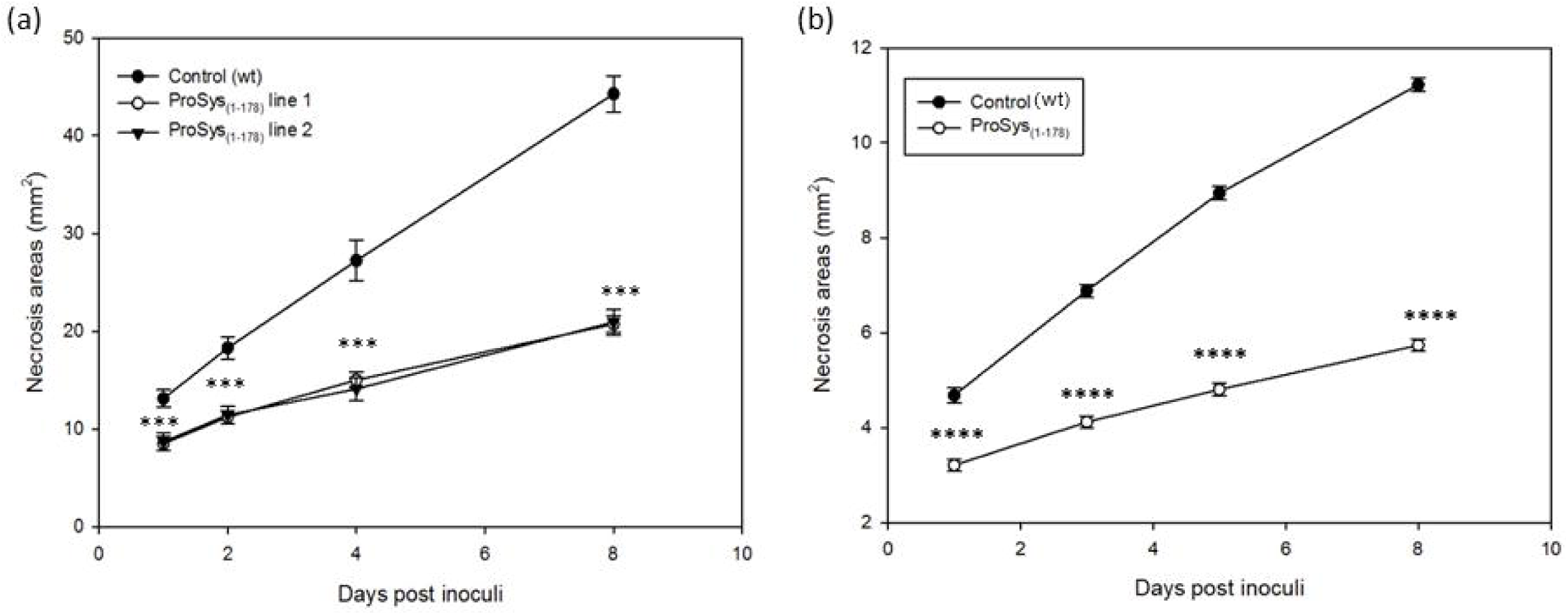
Enhanced resistance of ProSys_(1-178)_ transgenic lines and in ProSys_(1-178)_-treated tomato leaves to *B*.*cinerea*. (**a**) Dimension of necrosis areas in leaves of control plants and of ProSys_(1-178)_ transgenic lines 1 and 2. (**b**) Response to *B. cinerea* infection in leaves from mock-treated and ProSys_(1-178)_ treated plants. The graphs display the average (± S. E.) of the lesion size at several days post inoculum (hpi). Statistical analysis was performed with a Student’st-test (*** *P*< 0.001; **** *P*< 0.0001).Asterisks denote mean values that are significantly different.

### The transcriptomic profiles of tomato plants is strongly influenced by ProSys_(1-178)_ expression

The expression of the ProSys_(1-178)_ in tomato imposed a strong modification of the transcriptomic profile, up-regulating 428 and down-regulating 537 transcripts. The classification of the differentially expressed genes (DEGs), based on the ontological domain “biological process”, is shown in FigureS4. Several defense-related functional categories, such as “response to abiotic stimulus”, “response to biotic stimulus”, “secondary metabolic process”, and “cell death” were modified in transgenic plants. Tables S4 and S5 list all differentially expressed transcripts identified in ProSys_(1-178)_ plants. The identification of pathways including DEGs was carried out using KEGG analysis (Table S6). Defense-related pathways affected by ProSys_(1-178)_ expression were those involved in flavonoid biosynthesis and glutathione metabolism (Table S6).

Microarray data were validated by monitoring the expression of a group of genes by Real Time RT-PCR confirmingwhat observed through the array (FigureS5).

Tomato genes whose transcripts were modulated by the constitutive expression of the truncated Prosystemin were grouped according to their functional annotation (Table S4, S5).

#### Defense-related genes

A wide array of genes involved in early signaling responses were up-regulated, like four genes associated to the oxidative burst [i.e., glutathione-S-transferase (GST; Solyc09g011630.2.1, Solyc09g011500.2.1),NADH dehydrogenase (Solyc02g092270.2.1), laccase 22 (Solyc03g083900.2.1) and metacaspase 7 (Solyc09g098150.2.1)], two transcripts of onegenecoding for polygalacturonase (PG; Solyc08g082170.2.1)and a long list of transcripts encoding for kinases, phosphatases (including a dual specificity phosphatase 1, DUSP1; Solyc05g054700.2.1) and calcium related proteins. Other transcripts encoding for proteinsinvolved in early stages of defense responses were down-regulated, including members of the GST family (Solyc01g081250.2.1), peroxidases(Solyc01g104860.2.1, Solyc01g105070.2.1, Solyc07g017880.2.1, Solyc09g007270.2.1), catalases (Solyc12g094620.1.1) and calmodulin (Solyc03g115930.1.1, Solyc01g010020.2.1) (Table S5). A wide group of DEGs are associated with responses to abiotic stresses,like high temperature and include transcriptscoding several types of chaperone proteins (Solyc07g006540.2.1, Solyc05g055310.2.1), heat shock protein 4 (Solyc03g123540.2.1) and 70 (Solyc08g082820.2.1), stress-related protein (SRP; Solyc09g074930.2.1), DNAj heat shock proteins(Solyc03g117590.2.1, Solyc04g081530.1.1, Solyc05g053760.2.1, Solyc06g068500.2.1, Solyc08g005300.1.1) and dehydration-responsive family protein (Solyc04g080360.2.1).

#### Anatomical defensive structure

ProSys_(1-178)_ plants showed the up-regulation of genes involved in anatomical defensive structure such as those associated with the strengthening of physical barriers, like callose, cellulose synthases, and hydroxyproline-rich glycoprotein family, which participate in the formation of cell wall appositions designed to prevent or retard pathogen infiltration (Underwood, 2012). In addition, the strong up-regulation of transcripts encoding a PGenzyme (two transcripts related to the same gene (Solyc08g082170.2.1) and immune-responsive cytoskeletal elements, such as kinesin (Solyc02g084390.2.1, Solyc06g009780.2.1) and actin (one transcript, Solyc11g005330.1.1), further indicated the promotion of processes leading to cell wall reorganization.

#### Secondary metabolism

Plant antibiosis and antixenosis are often entrusted to secondary metabolism, which was altered in ProSys_(1-178)_ plants (FigureS4). The most representative secondary metabolism-related DEGs were found to be involved in flavonoid biosynthesis (i.e. crocetin, dihydroflavonol 4-reductase, chalcone synthase, flavanone 3 beta-hydroxylase). Another remarkable group of DEGs were involved in phenylpropanoid biosynthetic pathways.Interestingly, many phenylpropanoid show anti-microbial and anti-fungal activities (Naoumkina *et al*., 2010; Xu *et al*., 2011). In addition, the stress-related polyamines family was also affected, as shown by the up-regulation of the transcript of a putrescine-interacting protein (Solyc08g005860.2.1) and the down-regulation of an ornithine decarboxylase gene (Solyc04g082030.1.1), respectively.

#### Hormone-related pathways

Remarkably, numerous enzymes of the biosynthetic pathway of the three major plant hormones involved in defense responses, such as JA, salicylic acid (SA) and ethylene (ET), were down-regulated (Tables S4, S5). For example, transcripts involved in JA biosynthesis [i.e., 13-lipoxygenase (Solyc01g099210.2.1) and other classes of lipases]as well as members of the JA-responsive gene family [i.e., wound-inducible (Solyc03g093360.2.1), Kunitz trypsin inhibitor (Solyc03g098740.1.1), proteinase inhibitor I(Solyc09g084470.2.1) and metallocarboxypeptidase inhibitors (Solyc07g007250.2.1, Solyc07g007260.2.1)] were down-regulated. Similarly, transcripts involved in SA methylation, [i.e., S-adenosylmethionine-dependent methyltransferase(Solyc04g040180.2.1)] and SA-responsive genes [chitinase (Solyc01g097270.2.1), osmotin(Solyc08g080650.1.1), subtilisin (Solyc06g065370.2.1) and PR1a(Solyc09g006010.2.1) and PR1b (Solyc00g174340.1.1), those two latter may be also be responsive to ET] were also down-regulated (Leon-Reyes *et al*., 2009; Martínez-Medina *et al*., 2017; Molinari and Leonetti, 2019). A gene involved in ET biosynthesis, coding for 1-aminocyclopropane-1-carboxylate oxidase (Solyc09g089580.2.1), was also down-regulated, whereas other ET-related transcriptscoding for a serine/ threonine (Ser/Thr) kinaseand an ethylene receptor, were up-regulated. Ninegenes associated with the auxin pathway were down-regulated (Solyc01g110800.2.1, Solyc01g110940.2.1, Solyc02g077880.2.1, Solyc02g082450.2.1, Solyc03g082510.1.1, Solyc04g082830.2.1, Solyc05g008850.2.1, Solyc06g053260.1.1, Solyc10g052530.1.1). An analogous trend was observed for genes involved in gibberellin pathway as shown by the down-regulation of 6transcripts(Solyc05g051660.1.1, Solyc06g007890.2.1, Solyc07g056670.2.1, Solyc09g075670.1.1, Solyc11g017440.1.1, Solyc11g011210.1.1)involved in their biosynthesis and perception.

### Defense related genes are up-regulated in ProSys_(1-178)_ -treated plants

The effect of ProSys_(1-178)_ exogenous supply on tomato plants was evaluated by monitoring the expression of fivedefense related genes,selected among the genes up-regulated in transgenic plants and listed in Table S4, coding forpolygalacturonase (Solyc08g082170.2.1), dual specificity phosphatase 1(Solyc05g054700.2.1), basic leucine zipper protein family (Solyc01g090270.2.1), stress-related protein (Solyc09g074930.2.1)andglutathione S-transferase (Solyc09g011500.2.1). We included in this gene set *ProSys*(Solyc05g051750.2.1) to show it in comparison with the other genes.Transcript accumulation was analyzed 6 and 24 h after ProSys_(1-178)_ application. Notably, as shown in Figure5, all genes, except ProSys, were significantly over-expressed.

**Figure5:**
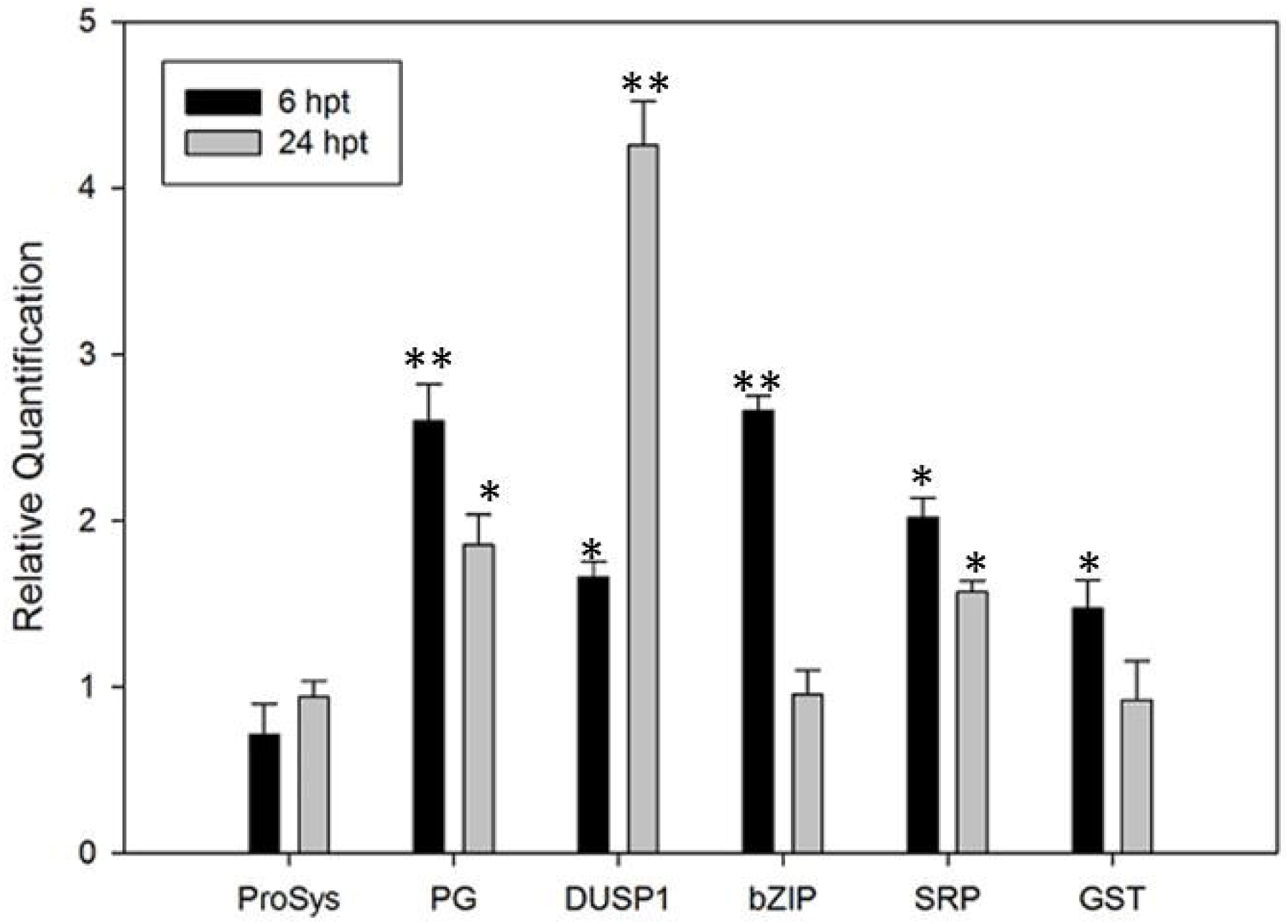
Relative expression of defense-related genes in tomato plants treated with exogenous ProSys_(1-178)_recombinant protein. Relative quantification of severaldefense-related genes induced at 6 h and 24 h after plant treatment (hpt) with 100 pMProSys_(1-178)_. Genes under investigation were: ProSystemin, ProSys; Polygalacturonase, PG; dual specificity phosphatase 1, DUSP1; basic leucine zipper protein family, bZIP; stress-related protein, SRP; glutathione S-transferase, GST.Quantities are relative to the calibrator represented by mock treated plants. Statistical analysis was performed with a Student’s t-test (* *P*< 0.05. ** *P*< 0.01).Asterisks denote mean values that are significantly different.

### ProSys_(1-178)_ induces the release of oligogalacturonides

In order to verify if the overexpression of PG observed both in transgenicProSys_(1-178)_and in ProSys_(1-178)_-treated plantswas associated with the release of OGs, we analyzed the leaves for OGs presence byMALDI-TOF analysis.

Figure S6 shows an example of MALDI-TOF spectra of the transgenic lines ProSys_(1-178)_ line 2 (back) and relative control (green).

We found signals attributable to OGs up to 4of degree of polymerization(DP) in transgenic plant samples and their relative quantification was obtained by comparing the peak intensity for each signal in the spectrum (Figure 6a). In all samples the OGs with 4 DP were the most abundant, especially in in ProSys_(1-178)_ line.

**Figure 6:**
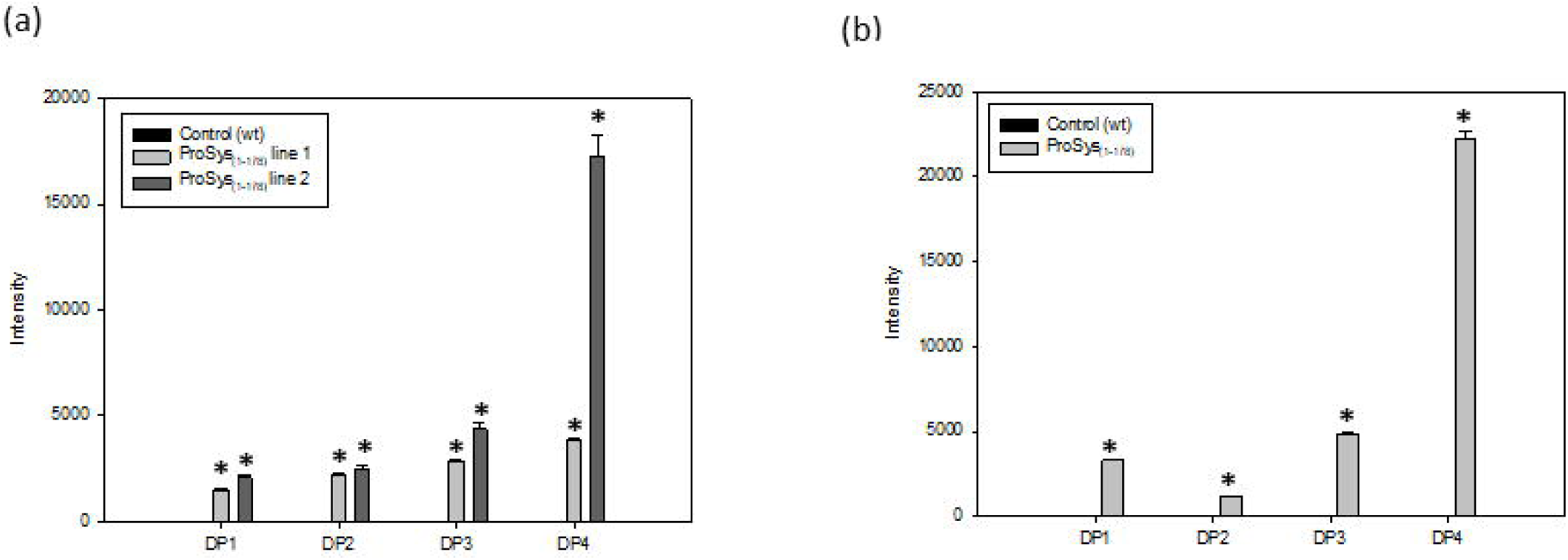
OGs identified in transgenic ProSys_(1-178)_ plants (a) and in ProSys_(1-178)_-treated plants (b). The graphs display means of different DP (± S. D.) of three biological replicates. DP = degree of polymerization.Statistical analysis was performed with a Student’s t-test (\**P*< 0.001).

Similarly in all ProSys_(1-178)_-treated samples, after 6 hour from the treatment,we found signals attributable to OGs with4 DP (Figure 6b).Relative quantification of OGs in the analysed samples are summarized in the graph below (Figure 6).

Interestingly both transgenic and treated plants released OGs similar in length.

## Discussion

Plants have evolved different families of functionally related peptide signals involved in defense responses against insects and pathogens. These peptides are often released by larger precursors and are perceived by membrane receptors, which activate a defensesignalling cascades(Albert, 2013; Farrokhi *et al*., 2008). One of the best characterized signaling peptides is tomato Sys which is released from ProSys upon wounding and herbivory. Sys interacts with aleucine-rich repeat receptor kinase(Wang *et al*., 2018)to trigger wound and defense responses in tomato(Corrado *et al*., 2007; Coppola *et al*., 2015; Coppola *et al*., 2019; Degenhardt *et al*., 2010; El Oirdi et al., 2011; McGurl *et al*., 1994; Pearce *et al*., 1991; Ryan, 2000; Zhang *et al*., 2020).

Despite these observations,recent findingsdescribingthe disordered structure of ProSys(Buonanno *et al*., 2018) suggested that, beside Sys, other ProSys regions couldhave roles in defense responses. Intrinsic disordered sequences allow proteins to bind multiple molecular partners with often different functional outcomes (Kim *et al*., 2008; Wallmann and Kesten, 2020). This mechanism mayexplain the ability of ProSysto protect plants against a wide array of stresses. This reasoning and previous work, showing proteomic reprogramming and enhanced resistance against fungi by tobacco plants expressingProSys_(1-178)_(Corrado *et al*., 2016), stimulated the hypothesis that ProSys is more than a simple precursor. Here we demonstrate that*S. littoralis* larvae fed onProSys_(1-178)_leaves had reducedgrowth and vitality, and that*B. cinerea* infection is strongly limitedon ProSys_(1-178)_plants. Interestingly, these results are confirmed by plant treatments with recombinantProSys_(1-178)_ protein. These findings are consistent with the deep transcriptomic reprogramming observed in transgenic plants constitutively expressing the deleted gene and with the expression profiles of defense-related genes evidenced in plants treated with recombinant ProSys_(1-178)_. As expected for the constitutive expression of a sequence associated with the plant immune system, metabolic and cellular processes were largely affected. The multilevel distribution of GO categories indicateda cellular reprogramming, involving primary and secondary metabolism, with a clear influence on defense mechanisms that included the up-regulation of early defense-related genes, like GST (Solyc09g011630.2.1, Solyc09g011500.2.1), associated with the regulation of the oxidative burst. Plant glutathione S-transferases are ubiquitous multifunctional enzymes encoded by large gene families that participate in ROS scavenging, stress tolerance,detoxification of toxic substances, plant growth and development, both *in vivo* and *in vitro*(Cairns *et al*., 2006; Gallé *et al*., 2019; Gulyás *etal*., 2014;Vernoux *et al*., 2000). Their fulfillment of such ansample number of functions mightexplain the observed up and down-regulation of different members of this gene family in the transgenic plants. Several other genes associated with the initial stages of the defense signaling cascade were up-regulated, such as kinases, phosphatases and calcium-related proteins. These genes are major milestones of defense response activation (Walling, 2000; 2009). For example, the expression of *DUSP1*(Solyc05g054700.2.1) is induced by cellular stresses and modulates selected MAP kinases, which are important plant defense components (Camps*et al*., 2000). Protein kinases and Ca^2+^ binding proteins play important roles in mediating defenseresponses against herbivores, while receptor-like kinases play a central role in pathogen recognition and the subsequent activation of plant defense mechanisms (Gao *et al*., 2014; Liu *et al*., 2016). Other increased transcripts code for members of the hydroxyproline-rich glycoprotein family (HRGPs; Solyc05g009930.2.1), a superfamily of cell wall proteins, involved in stress responses, signaling, and molecular recognition pathways (Johnson *et al*., 2017). Intriguingly, tomato HRGP (*Le*proHypSys), the protein precursor of HypSys I, II, and III peptides, that is up-regulated by wounding, Sys and methyl jasmonate treatments (Pearce and Ryan, 2003), and associated with Sys in the coordination of tomato defense responses (Narváez-Vásquez *et al*., 2007), has characteristic features of IDPs, similarly to ProSys(Johnson *et al*., 2017). Thus, these two IDPs may promote the establishment of a defense protein network active in protecting tomato plants against a wide array of biotic stressors. Interestingly, HRGPs appear to participate to cell wall appositions, to prevent or retard pathogen infiltration (Underwood, 2012). Indeed transcriptomic results suggest that ProSys_(1-178)_ highlypotentiates tomato physical barriers up-regulating a conspicuous group of genes related to cell wall reinforcement and callose synthesis. It was previously demonstrated that, the constitutive expression of callose synthase confers SA- and JA-independent resistance to powdery mildew in *Arabidopsis*(Ellinger and Voigt, 2014). Similarly, ProSys_(1-178)_ plants, in which SA and JA defense pathways are down regulated, showed the up-regulation of *Callose Synthase 11*(Solyc02g078230.1.1). According to these data the improved tolerance to *B. cinerea*shown by ProSys_(1-178)_ plantscould be associated with the strengthening of physical barriers, independently from hormone-regulated pathways. In this context, the abundance of transcripts coding for a PGenzyme(Solyc08g082170.2.1) likely led to the production of OGs, as observed both in transgenic and in treated leaves, that contributed to the observed resistance of ProSys_(1-178)_ plants against *B. cinerea*, as recently observed in *Arabidopsis*(Voxeur *et al*.,2019). The crucial role of the cytoskeleton and its re-organization during plant–pathogen interactions has been widely reported (Hardham *et al*., 2007; Hardham, 2013; Moral *et al*., 2017; Schmelzer, 2002). Thus, the up-regulation of genes coding for kinesin(Solyc02g084390.2.1, Solyc06g009780.2.1) actin (Solyc11g005330.1.1), and villin 2 (Solyc04g015830.2.1) and villin4 (Solyc02g021420.2.1) observed in ProSys_(1-178)_ plants suggests an extensive cytoskeleton re-organization. Especially in defense against fungi and oomycetes, actin dynamics represents one of the key-components in the formation of the physical barriers against their penetration (Janda *et al*., 2014). In addition to the reinforcement of physical barriers, the resistance of ProSys_(1-178)_ plants against insect pests could be the results of the modifications affecting secondary metabolism. The KEGG analysis helped to address 4 DEGs to the flavonoid biosynthesis pathway. These compounds are known to defend plants against various biotic and abiotic stresses including UV radiation, pathogens and insect pests (Shi *et al*., 2019). Among them is crocetin, an isoprenoid precursor of terpenoids, the largest group of plant chemicals with a primary role in plant growth and development, but also active in counteracting abiotic and biotic stressors (Tholl, 2015). Moreover, this molecule has been recently associated with plant-virus interactions (Parizad *et al*., 2019). Taken together, these observations explain the increased tolerance of the plants expressing ProSys_(1-178)_protein towards *B. cinerea* and *S. littoralis*, since cell wall reinforcement and phenylpropanoid-derived compounds are well recognized for their anti-feedant and cytotoxic effects on insect pests and pathogens (Bernards and Båstrup-Spohr, 2008).

Notably, transgenicplants showed the suppression of the major hormone-regulated pathways related to plant defense responses (i.e. JA, SA and ET), as indicated by the down-regulation of genes coding for lipoxygenase (LOX; Solyc01g099210.2.1),PIs (Solyc09g084470.2.1), pathogenesis-related proteins (PRs; Solyc09g006010.2.1, Solyc00g174340.1.1) and1-aminocyclopropane-1-carboxylate oxidase (*ACCO;* Solyc09g089580.2.1), a key enzyme involved in ET biosynthesis.

It is worth noting that ETperception occurs through hormone binding to receptors which act as negative regulators of ethylene response. When the hormone is absent ETreceptors activate proteins of the serine/threonine kinase family that act by phosphorylating a specific protein in order to repress its ability to induce ethylene responses (Gallie, 2015). In our results, genes coding for both (Ser/Thr) kinase(Solyc03g006770.2.1, Solyc06g075160.2.1, Solyc07g055860.2.1, Solyc08g013940.2.1, Solyc12g099320.1.1)and an ethylene receptorwere up-regulated coherently with the down-regulation of *ACCO*. In addition, genes involved in auxin (Solyc01g110800.2.1, Solyc01g110940.2.1, Solyc02g077880.2.1, Solyc02g082450.2.1, Solyc03g082510.1.1, Solyc04g082830.2.1, Solyc05g008850.2.1, Solyc06g053260.1.1, Solyc10g052530.1.1). and gibberellin pathways (Solyc05g051660.1.1, Solyc06g007890.2.1, Solyc07g056670.2.1, Solyc09g075670.1.1, Solyc11g017440.1.1, Solyc11g011210.1.1)were also down-regulated. Considering that the Sys signal transduction pathway, that leads to the production of JA, is activated by the interaction of Sys with its receptor (Wang *et al*., 2018), the lack of up-regulation of JA dependent genes in ProSys_(1-178)_ plants wasnot surprising since Sys was missing in the deleted genes that did not activate the endogenous *ProSys*. This gene is the major actor of the activation of genes involved in multiple hormone signaling pathways associated with plant defense(Coppola *et al*., 2015; Corrado *et al*., 2016; Orsini *et al*., 2010; Zhang *et al*., 2020). However, why the hormone related defense pathways are mainly down-regulated in ProSys_(1-178)_ plants needs further clarification. As mentioned, a remarkable up-regulation of PGrelated transcripts was observed in transgenic plants. These enzymes depolymerase pectin, one of the major plant cell wall components producing pectin derived OGs. OGs are perceived by cell wall-associated kinases and activate the plant innate immunity(Ferrari *et al*., 2013; Bacete *et al*., 2018). The down-regulation of the auxin biosynthetic pathways observed in ProSys_(1-178)_ plants is consistent with previous observation reporting that OGs have an antagonistic role to auxin (Savatin *et al*., 2011; Qi *et al*., 2012). This antagonism may play an important role in prioritizing defense*vs* growth thus allowing plants to allocate energy in the defense mechanisms if required. Interestingly, *PG* was up-regulated in plants treated with ProSys_(1-178)_ recombinant proteinconfirming that transcripts of this family are selectively up-regulated by ProSys_(1-178)_ protein since their expression profiles were not altered in the transcriptome of transgenic plants expressing prosystemin as analyzed through microarrays (Coppola *et al*., 2015). Therefore it appears that ProSys_(1-178)_ is not only biologically active but that it operates via a peculiar mechanism that is able to activate defense pathways that involve OGs. OGs are key components of DAMPsignaling able to elicit, in several plant species, a wide range of defense responses(Voxeur*et al*., 2019; Ferrari *et al*., 2013). They are thought to be released from plant cell walls upon partial degradation of homogalacturonan, originating during microbial infections,by microbial PGs (Cervone*et al*., 1989) or by the action of endogenous PGs induced by mechanical damage (Orozco-Cardenas and Ryan, 1999).Very recently it was shown that OGs are also produced by the activity of pectin lyases at least in Arabidopsis (Voxeur*et al*., 2019)

One structural requirement for the biological activity of OGs is the degree of polymerization. Although it has been suggested that long OGs (DP > 10) are the most effective in modulating signaling involved in plant defense responses (Denoux*et al*., 2008; Federici *et al*., 2006; Ferrari *et al*., 2007), it was shown that also short OGs (DP < 10) impact plant defense (Davidsson*et al*., 2017). For example, short OGs (DP4-6, DP2 and DP1-7, respectively) induced genes involved in pathogen response and defense in potato and tomato,while in tobacco and tomato they induced the synthesis of phytohormones (Montesano *et al*., 2001; Ridley *et al*., 2001; Simpson *et al*., 1998; Thain*et al*., 1990; Weber *et al*., 1996).Therefore we propose that the short OGs produced both in in ProSys_(1-178)_ transgenicand treated plants do actively contribute to defense responses.

Also remarkable wasthe up-regulation in ProSys_(1-178)_-treated plants of a member of *bZIP* transcription factor (TF) (Solyc01g090270.2.1)family, 6 h post plant treatment confirming the early involvement of bZIP TF in plant defense responses, in line with the key-roles played by TF in plant innate immunity (Li *et al*., 2016; Norman-Setterblad *et al*., 2000).

Collectively, the experimental evidences gathered so far clearly demonstrate that ProSys_(1-178)_ protein triggers tomato plant defense responses by activating specific classes of defense-related genes and prioritizing defense in respect to growth. It is tempting speculating that the overall function of the wild type prosystemin is to activate at least two tomato defense pathways, the Sys-dependent pathway connected with the induction of JA biosynthesis and the successive activation of a set of defense-related genes and the ProSys_(1-178)_-dependent pathway associated with OGs production leading to the OGs mediate plant immunity. Further studies are required to confirm this hypothesis. This presumed mechanism may explain the large ‘anti-stress’ capacity of ProSys. If and how this mechanism is dependent on the disordered structure of the precursor needs also further studies.

## Materials and Methods

### Plant material and growth condition

Tomato seeds (*Solanum lycopersicum* L. cultivar “Red Setter”) were surface-sterilized with 70% ethanol for 2 min, rinsed, washed with 2% sodium hypochlorite for 10 min and rinsed at least five times with sterile distilled water. Seeds were then germinated in petri dishes on wet sterile paper and kept in the dark for 3 days in a growth chamber at 24 ± 1°C and 60% of relative humidity (RH). Upon roots emergence, plantlets were transferred to a polystyrene tray with barren sterile S-type substrate (FloraGard; Oldenburg, Germany) in a growth chamber at 26 ± 1°C and 60% RH with a 18: 6 h light/dark photoperiod. After 2 weeks, plants were transferred to in 9 cm diameter pots filled with a sterile soil mixture using the same growth conditions.

### Tomato transgenic plants production and analysis

The pPRO binary vector (Corrado *et al*., 2016), containing the coding region of the tomato prosystemin cDNA lacking the last exon coding for the Sys peptide under the control of the CaMV 35S RNA promoter and the pea *rbcS* terminator, was used for the genetic transformation of *S. lycopersicum* L. “Red Setter” as previously described (Coppola *et al*., 2015). Putative transformants, selected on kanamycin (50 μg/ml), were analyzed by PCR to detect the transgene, as already described (Coppola *et al*., 2015). The isolation of total RNA from leaves of four-week old plants grown in sterile soil, the synthesis of the first strand cDNA and Real Time RT-PCR were performed as previously reported (Corrado *et al*., 2012). Transgenic plants are hereafter referred as ProSys_(1-178)_ plants. According to the transgene expression levels, 2 plants of T_0_ generation were reproduced up to T_4_generation to select genotypes homozygous for a single copy of the transgene. These lines are indicated as line 1 and 2.

### Molecular cloning, expression, and purification of ProSys_(1-178)_

ProSys_(1-178)_ was obtained after PCR amplification of prosytemin cDNA (GenBank: AAA34184.1) with site-specific synthetic primers (Table S1) and was cloned in the pETM11 protein expression vector. The generated plasmid was checked by DNA sequencing and appropriate digestion with restriction enzymes. The recombinant product was expressed in *E. coli*, BL21(DE3) strain, induced with 1 mM isopropyl-β-D-1-tiogalattopiranoside (IPTG) for 16 h at 22°C in 2-YT broth. Cells were harvested by centrifugation (20 min at 6000 × g at 4°C) and the final cell pellets were lysed in 20 mM Tris-HCl pH 8.0, 20 mM imidazole, 50 mM NaCl, pH 8.0, in presence of 1 mM dithiothreitol (DTT), 0.1 mM phenylmethanesulfonyl fluoride (PMSF), 5 mg/ml DNase I, 0.1 mg/ml lysozyme, 1 μg/ml aprotinin, 1 μg/ml leupeptin and 1 μg/ml pepstatin protease inhibitors (Sigma-Aldrich; Milan, Italy). Cells were then disrupted by sonication and after centrifugation (30 min at 30000 × g at 4°C) the supernatant (soluble fraction) was purified by FPLC on a 1 ml HisTrap FF column (GE Healthcare; Milan, Italy) by stepwise elution, according to manufacturer’s instruction. Fractions containing ProSys_(1-178)_ protein were dialyzed in PBS 1X (Phosphate buffer saline, 10 mM phosphate, 140 mM NaCl, 2.7 mM KCl, pH 7.4), 100 μM PMSF, 1 mM DTT pH 8.0 using a dialysis membrane with a molecular weight cut-off (MWCO) of 3500 Da for 16 h at 4°C. ProSys_(1-178)_ was finally purified in PBS 1X by size exclusion chromatography (SEC) on a Superdex 75 10/300 HP column (GE Healthcare; Milan, Italy) and purity assessed by 15% SDS-PAGE using Bio Rad Precision Plus Protein All Blue Standards (10-250 kDa) as molecular weight ladder (Bio-Rad; Hercules, CA, USA).

### LC-ESI-MS, circular dichroism and light scattering analyses

LC-ESI-MS and CD spectra were performed as previously described (Truppo *et al*., 2012; Buonanno *et al*., 2018) to confirm protein identity and behavior. SEC-MALS-QELS analysis of ProSys_(1-178)_ was performed at 0.5 ml/min in PBS 1X, 100 μM PMSF, 1 mM DTT pH 8.0 on a SEC 2000 column (Phenomenex; Torrance, CA, USA) linked to an FPLC ÄKTA coupled to a light scattering detector (mini-DAWN TREO, Wyatt Technology; Santa Barbara, CA, USA) and to a refractive index detector (Shodex RI-101; Showa Denko, Tokyo, Japan). Collected data were processed using the ASTRA 5.3.4.14 software (Wyatt Technologies Corporation).

### Plant treatments with ProSys_(1-178)_

Fifteen spots of 2 μl of 100 pMProSys_(1-178)_ solution were gently placed on the abaxial surface of fully expanded healthy leaves of four weeks-old plants (a mock treatment with buffer was used as control). Treated leaves were collected 6 and 24 h after ProSys_(1-178)_ application for molecular investigations and for bioassays unless otherwise indicated.

### OGs extraction by chelating agent

The OGs extraction protocol was a modified version of the protocol described in Pontiggia*et al*. (2015). For each sample, about 50 mg of crushed fresh leaves were re-suspended in 1 ml of 70% ethanol, centrifuged 15 min at 14000 rpm. The supernatant was discarded and the pellet was washed twice with a chloroform: methanol (1:1, vol/vol) mixture, vortexed, and centrifuged at 14000 rpm for 15 min. The supernatant was discarded, and the pellet was washed twice with acetone, centrifuged at 14000 rpm for 15 min and dried under vacuum. The pellet was re-suspended in 200 μl of ultrapure water and kept overnight at 4°C on a wheel. After centrifugation for 30 min at 14000 rpm, the supernatant was discarded and the pellet was re-suspended in 200 μl chelating agent solution (ChA= 50 mM EDTA dissolved in 1 M NaOH) and incubated overnight at 4°C on a wheel. After centrifugation for 30 min at 14000 rpm, the supernatant containing the chelating agent soluble fraction (ChASF) was recovered. Oligogalacturonides (OGs) contained in the ChASF were precipitated with 1 ml of 80% ethanol at -20°C overnight. Pellets obtained by centrifugation at 14000 rpm for 30 min were washed twice with 80% ethanol and dried under vacuum. OGs were dissolved in 100 μl ultrapure water and subjected to MALDI-TOF analysis.

### MALDI-TOF (Matrix Assisted Laser Desorption Ionization-Time of Flight) Mass Spectrometry

One microliter of extracted OGs were mixed with 4 μl of MALDI matrix (50mg/ml of PA/HPA/AC 18:1:1) and 1 μl of mix was spotted on the MALDI plat. Mass spectra were recorded on a 5800 plus MALDI TOF-TOF mass spectrometer (ABI SCIEX) equipped with a reflectronanalyzer and used in delayed extraction mode with 4000 Series Explorer v3.5 software. MALDI-MS data were acquired over a 100−3000 m/z mass range in the positive ion reflector mode. Each spectrum represents the sum of 400 laser pulses from randomly chosen spots per sample position. Three biological replicates for the two transgenic lines and for ProSys_(1-178)_-treated plants were used for the analysis.

### Bioassays

#### Herbivory by S. littoralis larvae

The impact of the experimental plants on *S. littoralis*larvae (Lepidoptera, Noctuidae) was assessed as previously described (Coppola *et al*., 2019), starting from a larval population reared on an artificial diet (Di Lelio *et al*., 2014) at 25 ± 1°C, 70 ± 5% RH under a 16: 8 h light/ dark photoperiod. The feeding bioassays were performed under the same environmental conditions, in polystyrene rearing trays (RT32W, Frontier Agricultural Sciences; Newark, Germany), bottom-lined with 3 ml of 1.5% agar (w/v) on which 150 newly hatched larvae were deposited, in groups of 50 individuals, on leaf disks and allowed to develop fully to the 2^nd^ instar. The rearing wells were closed by perforated plastic lids (RTCV4, Frontier Agricultural Sciences; Newark, Germany).

Soon after molting to the 3^rd^ instar, 32 larvae, for each experimental condition, were singly transferred into new trays prepared as above, and were daily offered fresh leaf disks of uniform size (initially of 1 cm^2^; later, disks of 2, 3, 4 and 5 cm^2^ were offered to meet the increasing nutritional needs of the larvae), obtained from sub-apical leaves of 4 weeks-old plants.

For the experiments performed with transgenic plants, larvae were fed with tomato leaf disks of 2 transgenic plant line 1 and 2 and of untransformed control plants, while for the experiments with plants treated with 100 pMProSys_(1-178)_, leaf discs of treated and control plants were daily supplied to larvae, 6 h after plant treatment.

The survival rate was daily assessed, while the larval weights were recorded every 5 days, for the ProSys_(1-178)_ transgenic plants experiments, and every two days, for the ProSys_(1-178)-_treated plants experiments. These parameters were recorded until pupation.

#### Infection by the necrotrophic fungus B. cinerea

For this bioassay, 4-weeks-old ProSys_(1-178)_ transgenic plants were inoculated with fungal spores as previously described (Corrado *et al*., 2005). Briefly, spores of *B. cinerea* were suspended in sterile distilled water, filtered through sterile Kimwipes (Kimberly-Clark; Dallas, TX, USA) to remove hyphae fragments and adjusted to a concentration of 1×10^6^ spores/ ml. Ten μl of the spore suspension were applied between the leaf veins, using four different inoculation points per leaf. Five plants for each line 1 and 2 were used. Lesions diameters were measured at different time points (1, 3, 5 and 8 days post inoculum) using a digital caliper (Neiko 01407A; Neiko Tools, Taiwan, China). For each sample 2 technical replicates were used.

The assay was also performed on detached leaves exogenous supplied with ProSys_(1-178)._ For this purpose, leaves of four weeks old plants were harvested and treated with 100 pM recombinant protein or control buffer and, after 6 hours, a spore suspension of *B. cinerea* was applied. Necrosis were measured as reported above. Three leaves from three different plants per each treatment were used.

### Two-Color Microarray-Based Gene Expression Analysis

Total RNA was extracted from leaves using the Plant RNeasy mini kit (Qiagen; Hilden, Germany), according to manufacturer’s protocol. RNA quantification and quality control were carried out with the 2100 Bioanalyzer system (Agilent Technologies: Santa Clara, CA, USA). Samples with a 260/280 nm absorbance ratio > 1.8 and a 260/230 nm absorbance ratio > 2 were labeled and hybridized to the Tomato Gene Expression Microarray 4 × 44K (Agilent Technologies) as previously described (Coppola *et al*., 2015). Experiments were run in triplicate for each biological condition. Upon scanning and image data processing, raw data were analyzed using Genespring GX 10 software (Agilent Technologies). Statistical analysis was performed using background-corrected mean signal intensities from each dye channel. Microarray data were normalized using intensity-dependent global normalization (LOWESS). Differentially expressed RNAs were identified after filtering by the Benjamini and Hochberg False Discovery Rate (*P*< 0.05) and a minimum of a 2-fold variation in expression compared to untransformed controls. Differentially expressed genes (DEGs) functional annotation was carried out by sequence analysis using the Blast2GO software (Götz *et al*., 2008). Mapping of enzymatic activities into molecular pathways was acquired from the Kyoto Encyclopedia Gene and Genomes (KEGG) database.

Microarray data were validated by Real Time RT-PCR targeting 7 DEGs. The expression analysis was extended to other 2 transgenic genotypes (ProSys_(1-178)_ line 1 and 2).

### Relative quantification of gene expression

Expression analysis were carried out by Real Time RT-PCR using Rotor Gene 6000 (Corbett Research; Sydney, Australia). The isolation of total RNA from leaves of four-week old plants grown in sterile soil, the synthesis of the first strand cDNA and Real Time RT-PCR were performed as previously reported (Corrado*et al*., 2012).Two technical replicates for each of the three biological replicates per samples were used. The housekeeping gene *EF-1α* was used as endogenous reference gene for the normalization of the expression levels of the target genes. Relative quantification of gene expression was carried out using the 2^-ΔΔCt^ method (Livak and Schmittgen, 2001). Primers and their main features are described in Table S1.

### Statistical analysis

Relative quantification of transcript abundance was compared by Student’s t-test when tests were compared to controls. In other cases, when multiple comparisons were considered, One-Way ANOVA was applied.

Survival curves of *S. littoralis* larvae fed with experimental tomato leaf were compared by using Kaplan-Meier and log-rank analysis. Normality of data was checked with Shapiro-Wilk test and Kolmogorov-Smirnov test, while homoscedasticity was tested with Levene’s test and Barlett’s test. Unpaired Student’st-test or One-Way ANOVA test, followed by Tukey’s post-hoc multiple-comparison test, were used to compare larval weight, respectively for bioassay with tomato treated plants and tomato ProSys_(1-178)_ transgenic plants. When necessary, transformation of data was carried out to meet the assumptions of normality and homoscedasticity. When significant effects were observed (*P*< 0.05), the Tukey’spost-hoc test was used to compare mean values. Error bars referring to deviation standard were displayed.

Student’s t-test was also used to compare necrosis area for bioassay with tomato treated plants and tomato ProSys_(1-178)_ transgenic plants. Moreover, the relative quantification of OGs abundance was compared by Student’s t-test between experimental samples and controls. Error bars referring to standard error were displayed.

## Acknowledgements

We would like to thank Materias S.r.l. for the encouragement and interest.This work was supported by the European Union’s Horizon 2020 research and innovation program, under grant agreement no. 773554 (EcoStack), by the project PGR00963 financed by the Italian Ministry of Foreign Affairs and International Cooperation and by the program STAR (Line 1, 2018) “Exitbat-EXploiting the InTeraction of Belo- and Above-ground plant biostimulants promoting the sustainable protection of Tomato crop” financially supported by University of Naples Federico II and foundation Compagnia di San Paolo, Italy.

## Author Contributions

DM and MC should be considered joint first author; DM produced and structurally characterized ProSys_(1-178)_, performed gene expression studies and contributed to manuscript writing; MC produced and analyzed transgenic plants, performed and analyzed microarray analyses, and contributed to manuscript writing;MB purified ProSys_(1-178)_; IDL and AB carried out insect bioassays and data analysis; SMM supervised the biochemical work; CM performed OGs extraction and MALDI-TOF mass spectrometry analyses; AA MALDI-TOF mass spectrometry analysis; G.C. contributed to microarray analysis; JDF participated to experimental work and revised the manuscript; FP contributed to insect work supervision and manuscript revision; RR conceived and supervised the work and wrote the manuscript.

## Supporting information

**Figure S1:Molecular analysis of ProSys**_**(1-178)**_ **transgenic plants**.(**a**) Schematic representation of the pPRO8 T-DNA binary vector used for plant genetic transformation. LB: T-DNA left border sequence; 35S: CaMV35S RNA gene promoter; ProSys_(1-178)_: prosystemin cDNA sequence lacking the Sys coding region; *rbcS*: terminator of the pea rubisco small subunit; *nptII*: neomycin phosphotransferase II coding sequence; *nos*: nopaline synthase promoter; RB: T-DNA right border sequence. (**b**) PCR screening of transgenic plants. Lane 1: 1 kb Plus ladder(Thermo Fisher Scientific); number on the left margin indicate the expected marker fragment size. in base pairs; lane 2: control (no template); lane 3: pPRO8 binary vector; lanes 4-10: amplification products from DNA isolated from putative transgenic plants. (**c**) Relative quantification (RQ) of ProSys_(1-178)_ expression (by real-time RT-PCR). Numbers indicate the genotype corresponding to single transformation events. Primers anneal on ProSys N-terminal region allowing the amplification on both transgenic and untransformed plants. RQ is relative to the calibrator genotype represented by the cDNA from the tomato ‘Red Setter’ cultivar. Statistical analysis was performed with a Student’s t-test(* *P*< 0.05; ** *P*< 0.01). Asterisks denote mean values that are significantly different.

**Figure S2:Features of purified recombinant ProSys**_**(1-178)**_. (**a**) 15% SDS-PAGE visualized by Coomassie Brilliant Blue. M: molecular mass marker (20-250 kDa); 1: purified ProSys_(1-178)_. (**b**) SEC calibration curve built using five standard proteins (red crosses); data for ProSys_(1-178)_ (full circle) were calculated from this curve; (**c**) SEC-MALS of ProSys_(1-178)_ confirmed that the protein is a monomer in solution.

**Figure S3:Structural changes in ProSys**_**(1-178)**_ **analyzed by far-UV circular dichroism CD**. (**a**) CD spectra at different temperatures (10°C, 20°C, 30°C, 40°C, 50°C and 60°C) recorded from 260 nm to 190 nm at 20°C in 10 mM phosphate buffer, pH 7.4. The ellipticity at 222 nm vs. temperature is shown in the inset. (**b**) CD spectra in the presence of increasing TFE concentration (0%. 15%. 20%. 25%, 30% and 35%) recorded from 260 nm to 190 nm at 20°C. The ellipticity at 222 nm vs TFE percentage is shown in the inset.

**FigureS4: Multilevel distribution of GO categories associated to DEGs identified in ProSys**_**(1-178)**_ **plants**. GO terms associated to up-regulated (red bars) and down-regulated (green bars) genes are based on the “Biological Process” ontological domain.

**Figure S5:Microarray validation by Real Time RT-PCR. Relative quantification of DEGs identified through microarray analysis**. Genes under investigation were: 3-oxo-5-alpha-steroid 4-dehydrogenase family protein, PPR2; MLO-like protein 3, MLO3; Crocetin UDP-glucosyltransferase family 1 protein, crocetin; stress-related protein, SRP; dual specificity phosphatase, DUSP1; 1 leucine aminopeptidase 1A. Lap1A; mate efflux carrier protein, MATE. Data were calibrated on the ‘Red Setter’ untransformed genotype. Statistical analysis was performed with a Student’s t-test (* *P*< 0.05; ** *P*< 0.01).

**Figure S6:OGs identified in ProSys**_**(1-178)**_**plants**.MALDI-TOF spectra of the transgenic lines ProSys_(1-178)_ line 2 (back) and its relative control RS1 (green). DP = degree of polymerization.

**Table S1**. Primer and their main features.

**Table S2**.Weight increase of *S. littoralis* larvae fed on leaves of ProSys_(1-178)_ transgenic plants. The leaves employed were taken from control plants (C) and from two transgenic plant lines: ProSys_(1-178)_ line 1 and 2, respectively. Average values ± standard deviations are reported for each sample for all experimental time points analyzed. For each time point, mean values denoted with different letters are significantly different (*One-way ANOVA: *P*< 0.0001 followed by Tukey’s post-hoc multiple-comparison test).

**Table S3**.Weight increase of *S. littoralis* larvae fed on leaves of tomato plants treated withProSys_(1-178)_recombinant protein. Averages ± standard deviations are reported for ProSys_(1-178)_-treated and control (C) samples for all experimental time points. (Student’st-test: *\* P* <0.01; *\*\* P* <0.0001).

**Table S4**. Differentially expressed genes (DEGs) of ProSys_(1-178)_ transgenic plants identified by microarray analysis (Tomato array 4×44K, Agilent Technologies). ProbeName, Locus ITAG2.5, Fold change, Best Blast Hit Descriptor for up-regulated genes are reported.

**Table S5**. Differentially expressed genes (DEGs) of ProSys_(1-178)_ transgenic plants identified by microarray analysis (Tomato array 4×44K, Agilent Technologies). ProbeName, Locus ITAG2.5, Fold change, Best Blast Hit Descriptor for up-regulated genes are reported.

**Table S6**.KEGG analysis of the differentially expressed genes.Mapping of enzymatic activities (Enzyme and Enzyme ID) of the DEGs (Sequence ID) in the KEGG pathways (Pathway).

