## Supplementary figures and images for "Tomato Prosystemin is much more than a simple Systemin precursor"

### Figure S1

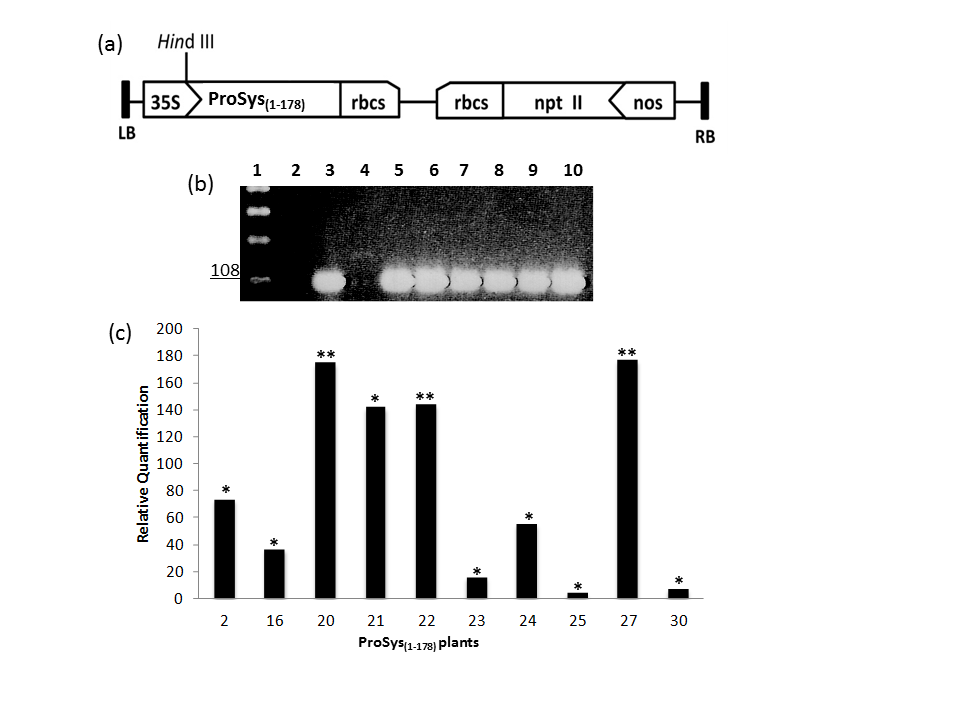
