## Supplementary Table S2 for "Tomato Prosystemin is much more than a simple Systemin precursor"

**Table S2.**Weight increase of *S. littoralis* larvae fed on leaves of ProSys_(1-178)_ transgenic plants. The leaves employed were taken from control plants (C) and from two transgenic plant lines: ProSys_(1-178)_ line 1 and 2, respectively. Average values ± standard deviations are reported for each sample for all experimental time points analyzed. For each time point, mean values denoted with different letters are significantly different (*One-way ANOVA: *P*< 0.0001 followed by Tukey’s post-hoc multiple-comparison test).

| Days of feeding | C  Larval weight (g) | ProSys_(1-178)_ line 1  Larval weight (g) | ProSys_(1-178)_ line 2  Larval weight (g) | F | P value |
| --- | --- | --- | --- | --- | --- |
| 1 | **0.0047** ± 0.0007 ^a^ | **0.0046** ± 0.0006 ^a^ | **0.0046** ± 0.0007 ^a^ | F (3. 124) = 1.584 | 0.1966 |
| 5 | **0.0144** ± 0.0037^a^ | 0.0131 ± 0.0027 ^a^ | **0.0101** ± 0.0016 ^b^ | F (3. 115) = 21.96 | 0.0001 * |
| 10 | **0.0226** ± 0.0091 ^a^ | **0.01642** ± 0.0054 ^b^ | **0.0159** ± 0.0039 ^b^ | F (3. 109) = 9.184 | 0.0001 * |
| 15 | **0.0964** ± 0.0341 ^a^ | **0.0266** ± 0.009 ^b^ | **0.0336** ± 0.036 ^b^ | F (3. 108) = 98.39 | 0.0001 * |
| 20 | **0.2763** ± 0.0913^a^ | **0.0545** ± 0.0134 ^b^ | **0.0455**± 0.0195^b^ | F (3. 98) = 144.1 | 0.0001 * |
| 25 | **0.9309** ± 0.0155 ^a^ | **0.08124** ± 0.0256 ^b^ | **0.0746** ± 0.0253 ^b^ | F (3. 91) = 2242 | 0.0001 * |
