## Supplementary Table S3 for "Tomato Prosystemin is much more than a simple Systemin precursor"

**Table S3.**Weight increase of *S. littoralis* larvae fed on leaves of tomato plants treated withProSys_(1-178)_recombinant protein. Averages ± standard deviations are reported for ProSys_(1-178)_-treated and control (C) samples for all experimental time points. (Student’st-test: ** P <*0.01; *** P <*0.0001).

| Days of feeding | C  Larval weight (g) | ProSys_(1-178)_  Larval weight (g) | t | dF | *P*value |
| --- | --- | --- | --- | --- | --- |
| 1 | **0.0039** ± 0.0005 | **0.0039** ± 0.0006 | 0.3865 | 62 | 0.7005 |
| 3 | **0.0082** ± 0.0026 | **0.004**8 ± 0.0015 | 6.373 | 62 | 0.0001 ** |
| 5 | **0.0156** ± 0.0044 | **0.0096** ± 0.0168 | 2.929 | 60 | 0.0599 |
| 7 | **0.0192** ± 0.0089 | **0.0084** ± 0.0029 | 5.918 | 56 | 0.0001 ** |
| 10 | **0.0321** ± 0.0083 | **0.0173** ± 0.0082 | 6.148 | 48 | 0.0001 ** |
| 12 | **0.0556** ± 0.0121 | **0.0143** ± 0.0067 | 11.87 | 42 | 0.0001 ** |
| 14 | **0.0849**± 0.0321 | **0.0455** ± 0.0587 | 2.926 | 28 | 0.0067* |
