## Supplementary Table S1 for "Tomato Prosystemin is much more than a simple Systemin precursor"

Supplementary Table S1. Primer and their main features. Table reports primer name, sequence, amplicon length, name/gene symbol and identifier (ITAG annotation) used in the experimental procedures.

| **Primer** | **Sequence (5’🡪3’)** | **Amplicon length (bp)** | **Name/Gene synbol** | **Accession Number** |
| --- | --- | --- | --- | --- |
| GST Fw | AGATGAGACATTTGAAGGCCCT | 106 | GST | Solyc09g011500.2.1 |
| GST Rv | TACCACCGCTCCCACCTTAT |  |  |  |
| EF Fw Rt | CTCCATTGGGTCGTTTTGCT | 101 | elongation Factor 1 alpha | Solyc06g005060.2 |
| EF Rv Rt | GGTCACCTTGGCACCAGTTG |  |  |  |
| bZIP Fw | CGGCGTGTGGAAGATGAGAT | 145 | bZIP | Solyc01g090270.2.1 |
| bZIP Rv | CCTCAAGGGCCTCTGTCATC |  |  |  |
| PPR2 Fw | ATACCCAGAACGACCCGGTA | 103 | PPR2 | Solyc01g110460.2.1 |
| PPR2 Rv | CACCACCAGTCCCGATTCTC |  |  |  |
| MLO3 Fw | CGGCACATTTAGCACCACAG | 136 | MLO3 | Solyc06g010030.2.1 |
| MLO3 Rv | GAGTAAGAAGAGCACGGCGA |  |  |  |
| Crocetin Fw | GGAGTACCTGTCGTGGCTTT | 83 | Crocetin | Solyc12g098590.1.1 |
| Crocetin Rv | ACTCCACTCTTCCACACATCT |  |  |  |
| SRP Fw | CTTAGTGGACACAGCGAGCA | 123 | Stress-related protein | Solyc09g074930.2.1 |
| SRP Rv | GATCTCCATGCCAAAACCGC |  |  |  |
| DSP Fw | TCATCAGCCTCATCACCTCC | 148 | Dual specificity phosphatase 1 | Solyc05g054700.2.1 |
| DSP Rv | TGGAGTCAGACGGTGTTGAA |  |  |  |
| Lap Fw | ATCTCAGGTTTCCTGGTGGAAGGA | 99 | leucine amino peptidase | Solyc00g187050.2 |
| Lap Rv | AGTTGCTATGGCAGAGGCAGAG |  |  |  |
| Mate Fw | ACCCATCAATGACACCCAAG | 154 | mate efflux protein | BI933305 |
| Mate Rv | GGCATGTGGTATGGGATGTT |  |  |  |
| PG Fw | TCAGCCCTTTTCCTTGTGCA | 125 | Polygalacturonase | Solyc08g082170.2.1 |
| PG Rv | TGCGACCACCATCATTTCCA |  |  |  |
| PcSysfw | TCTGAATTTGTCTCCCGTTAGAA | 119 | Endogenous Prosystemin | Solyc05g051750.2.1 |
| PcSysrv | AGCCAAAAGAAAGGAAGCAATAC |  |  |  |
| ProSys Fw | GGGAGGGTGCACTAGAAATA | 161 | ProSystemin | M61914 |
| ProSys Rv | TTGTCGAAACCGATGATACG |  |  |  |
